# Statistical analysis reveals the onset of synchrony in sparse swarms of *Photinus knulli* fireflies

**DOI:** 10.1101/2022.01.05.475109

**Authors:** Raphaël Sarfati, Laura Gaudette, Joseph M. Cicero, Orit Peleg

## Abstract

Flash synchrony within firefly swarms is an elegant but elusive manifestation of collective animal behaviour. It has been observed, and sometimes demonstrated, in a few populations across the world, but exactly which species are capable of large-scale synchronization remains unclear, especially in low-density swarms. The underlying question which we address here is: how does one qualify a collective flashing display as synchronous, given that the only information available is the time and location of flashes? We propose different statistical approaches and apply them to high-resolution stereoscopic video recordings of the collective flashing of *Photinus knulli* fireflies, hence establishing the occurrence of synchrony in this species. These results substantiate detailed visual observations published in the early 1980s and made at the same experimental site: Peña Blanca Canyon, Coronado National Forest, Arizona, USA. We also remark that *P. knulli*’s collective flashing patterns mirror that observed in *Photinus carolinus* fireflies in the Eastern United States, consisting of synchronous flashes in periodic bursts with rapid accretion and quick decay.

## 1 Introduction

Many animal species are capable of, and benefit from, behaving collectively, from insects forming rigid aggregates, such as ants and bees, to large mammals migrating as herds over thousands of kilometers. Although in some cases the emergence of collective dynamics is intuitively evident, for example in collective turns of flocking birds or swirling schools of fish, in other instances characterizing the ensemble structure or dynamics as collective requires a much finer analysis than simple visual observations. This is the case, for example, in disorganized midge swarms [1]. In fact, despite a growing interest in large-scale patterns of animal groups, a clear definition of which behaviour qualifies as collective is still lacking.

To address this broad and complex question, it may be easier to start with a simple subset of collective behaviour: synchrony. Animal synchronization manifests itself in many different ways and across various timescales [2], and it is certainly a signature of how social interactions produce system-wide patterns. An inspiring and readily accessible example of biological synchrony is seen sometimes on summer nights in firefly swarms, when most flashes occur at specific times.

Initially observed in Southeast Asia, synchronous fireflies were first reported in North America in the 1910s [3]. Observations were rare and sporadic, and often received with skepticism [4], precisely because rigorous demonstration of synchrony is difficult, especially in the absence of experimental data. In 1968, Buck provided photometric evidence to demonstrate the occurrence of synchrony in *Pteroptyx malaccae* congregations in Thailand [5]. In 1983, Cicero published a thorough account of lek behaviour in the Arizona firefly *Photinus knulli*, following extensive observations made by eye [6]. These observations did include the occurrence of synchrony. Beginning in the early 1990s, Eastern US synchronizing counterparts *Photinus carolinus* and *Photuris frontalis* have received considerable attention from scientists [7], bringing videographic evidence to characterize the occurrence [8, 9], mechanisms [10, 11], possible function [12, 13], and modalities of synchronous patterns [14]. In parallel, this renewed interest in firefly synchronization motivated numerous developments of mathematical and computational models [15, 16, 17].

In these experimental studies, thanks to the very high number of fireflies flashing in unison, the occurrence of synchrony was simply accepted from observations and raw data, without demanding further statistical analysis. The high signal-to-noise ratio was sufficient proof. Since then, recurrent speculation about possible synchrony in other species illustrates the need for a general methodology, adapted in particular to low-density populations.

This paper proposes to address the following question: How does one characterize synchronization behaviour amongst several dispersed fireflies? The situation at hand is different from other typical situations where synchrony is involved. In mathematical models and numerical simulations, where each agent’s internal phase *θ_k_*(*t*) is known at all times, it is possible to calculate an average phase order parameter, *ψ*(*t*) = arg (〈*e^iθ_k_^*)*_k_*), whose value between 0 and 1 quantifies the degree of synchrony within the system [18]. In situations where only firing is detectable, but each agent is continuously tractable, such as systems of neurons, it is possible to calculate cross-correlations between firing times of different constituents [19]. For fireflies, however, their internal phase is unknown, and in their natural habitat individual fireflies cannot be tracked for more than the duration of a flash train, after which they vanish in the obscurity. We propose here two approaches to demonstrate synchrony from high-resolution video-recordings: 1) from the time series of number of flashes in a camera’s field-of-view; and 2) from the spatiotemporal correlations between flash occurrences, after 3D reconstruction of the swarm.

## 2 Methods

The general experimental area was the same as described in detail in Ref. [6], namely the Peña Blanca Canyon (above lake) within the Coronado National Forest (Pajarito Mountains, Santa Cruz County, Arizona, USA). It comprised an intermittent river bed with a gravel road and campground nearby. The area where video recordings occurred was situated on the opposite side of the canyon from the road and covered with dense vegetation (Fig. 1A).

**Figure 1:**
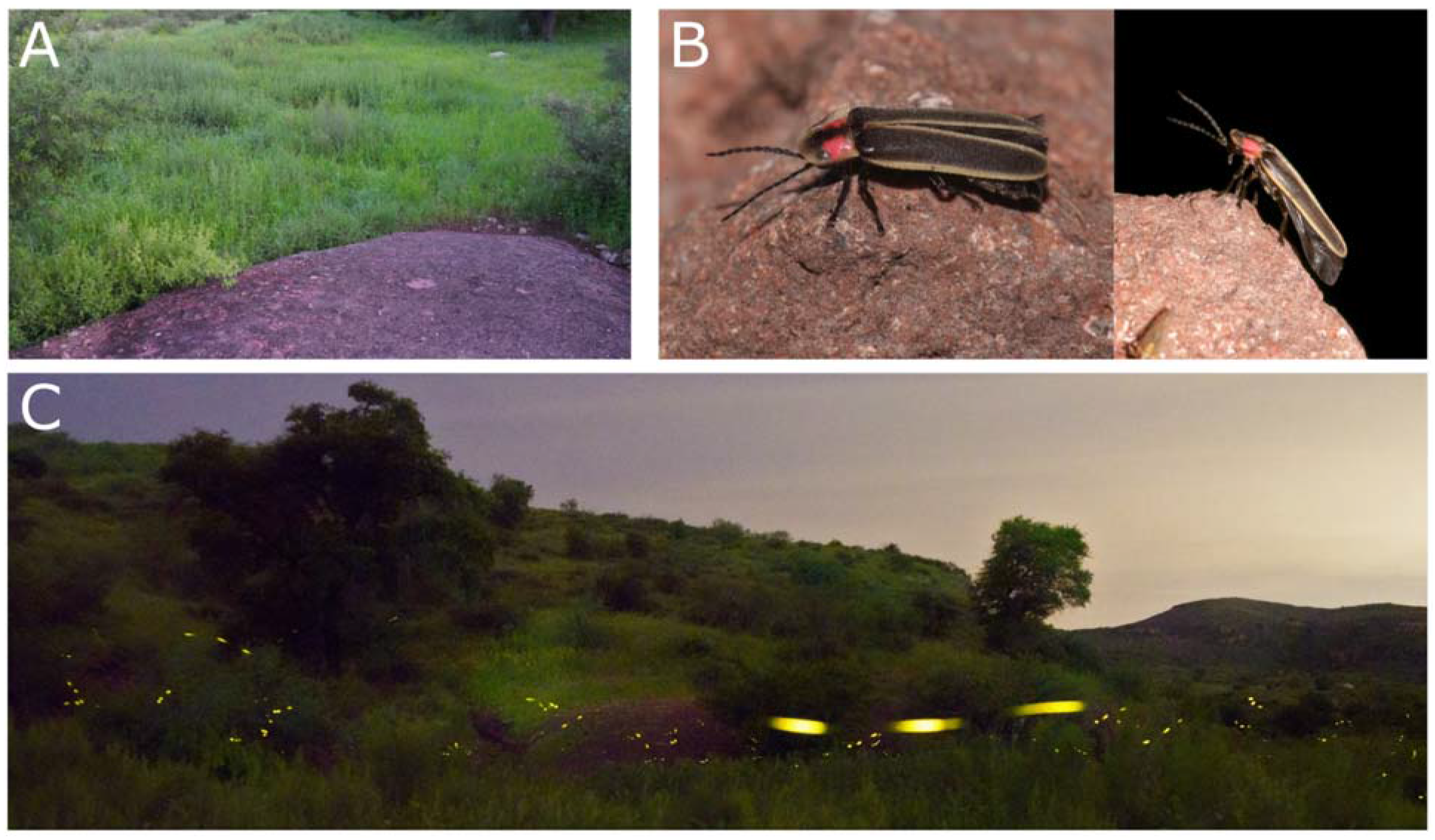
(A) Field-of-view from one of two recording cameras, near the river bed of the Peña Blanca Canyon. (B) *P. knulli* male firefly. (C) Long exposure (15s) photograph of collective display of *P. knulli* near the recording site. Several flash triplets are apparent.

Field observations occurred every night between 4 August and 13 August, 2021. The first flashes could generally be seen between 7:50-8pm, local time (Mountain Standard Time, MST). Sunset occurred around 7:15pm MST. Due to initial adjustments and varying environmental conditions (rain, transient flooding), three full datasets of stereoscopic recordings were eventually obtained (August 7, 9, 10, about 150min each). Recordings were obtained as described previously (see Ref. [14]) from Sony α7R4 cameras at 60 frames-per-second (fps) mounted with a wide-angle lens. Video processing and data analysis were performed with Matlab. Flash centroids were extracted by intensity thresholding after background subtraction. For stereoscopic recordings, cameras pairs were calibrated using about 10 pairs of pictures of a checkerboard (25cm side length) and the stereoCalibration toolbox. Flash 3D positions were subsequently obtained by triangulation.

In order to confirm species identification, a few specimens were collected every night with an insect net and carefully inspected, then released. A few specimens were photographed as well.

## 3 Results

### 3.1 Preliminary observations

#### Species identification

Collected male individuals (Fig. 1B), either on the ground or in-flight, were all consistent with previous descriptions of *P. knulli* morphology [6]. From collected specimens and general visual observations, it appeared that only one species was displaying (flashing) over the course of our field experiments. Of note, however, glowing larvae, identified based on direct rearing as possibly *Pleotomus nigripennis* LeConte, were also observed in the leaf litter and on tree trunks.

#### Individual flashing

By eye and in close proximity, *P. knulli* flying males appear to emit flash triplets spanning a period of about 1s (Fig. 1C). Occasionally, flash phrases of two, four, or five flashes can also be observed, but are much less common. Flash triplets from a single individual are typically separated by at least 3-5s. Males were dispersed and solitary in their emergence.

### 3.2 Time series reveal spikes of correlated activity

To analyse patterns of collective flashing in *P. knulli*, we first calculated the number *N* of flashes detected in each movie frame (Fig. 2A). Since firefly activity was relatively low, most frames contained no flash, some captured one single flash, and a few captured between two and five flashes, under the given experimental conditions. While synchrony implies that several flashes occur at the same time, the reverse proposition is not necessarily true: concurrent flashes could happen by accident, even with no underlying interactions between fireflies. Intuitively, however, if the proportion of concurrent flashes is large, it tends to indicate intrinsic correlations. To quantitatively evaluate this hypothesis, we must compare the observed distribution of flashes *P*(*N*) against the null hypothesis that flashes happen independently at random. Independent events happening within a given time frame at a constant rate λ are described by the Poisson distribution: *Poiss*(*N* = *k*) = λ*^k^e*^−λ^/*k*!. In our situation, because the swarm’s flashing rate was fairly constant around mean value λ = 0.099 flashes/frame (see SI, Fig. S3), we can compare the experimental distribution *P*(*N*) with the Poisson distribution with the same average, as shown in Fig. 2B. Evidently, the proportion of large values of *N* is vastly superior (about one order of magnitude) to what would be predicted from the Poisson distribution, hinting that concurrent flashes are too frequent to occur only by chance. To quantify this discrepancy, we employed goodness-of-fit (gof) tests for the Poisson distribution. Several such tests have been developed [20]. For their simplicity of implementation and interpretation, we used two test statistics, *Z* and *T*, based respectively on the first and second, and third and fourth, moments of the empirical distributions [20, 21]. As a brief reminder, the Poisson distribution has mean equal to variance, and squared-skewness equal to excess kurtosis. Significant deviations from these equalities indicate non-Poissonity. Thanks to appropriate normalizations, both *Z* and *T* are asymptotically distributed according to a normal distribution of mean 0 and variance 1 (see SI, Fig. S1). For the empirical distribution of *N* in Fig. 2B, we find *Z* = 55 and *T* = −28, which significantly deviate from the range of values expected if the underlying distribution were Poisson (corresponding *p*-values are *p_Z_* ~ 10^−67^ and *p_T_* ~ 10^−18^, see SI Section 3). Therefore, the null hypothesis is rejected by the test statistics, at a very high significance level. Collective flashes are not independent, and *P. knulli*’s collective display can be considered synchronous with extremely high probability, at least episodically. Similar conclusions can be drawn from the time series from the two other datasets (see SI, Fig. S3).

**Figure 2:**
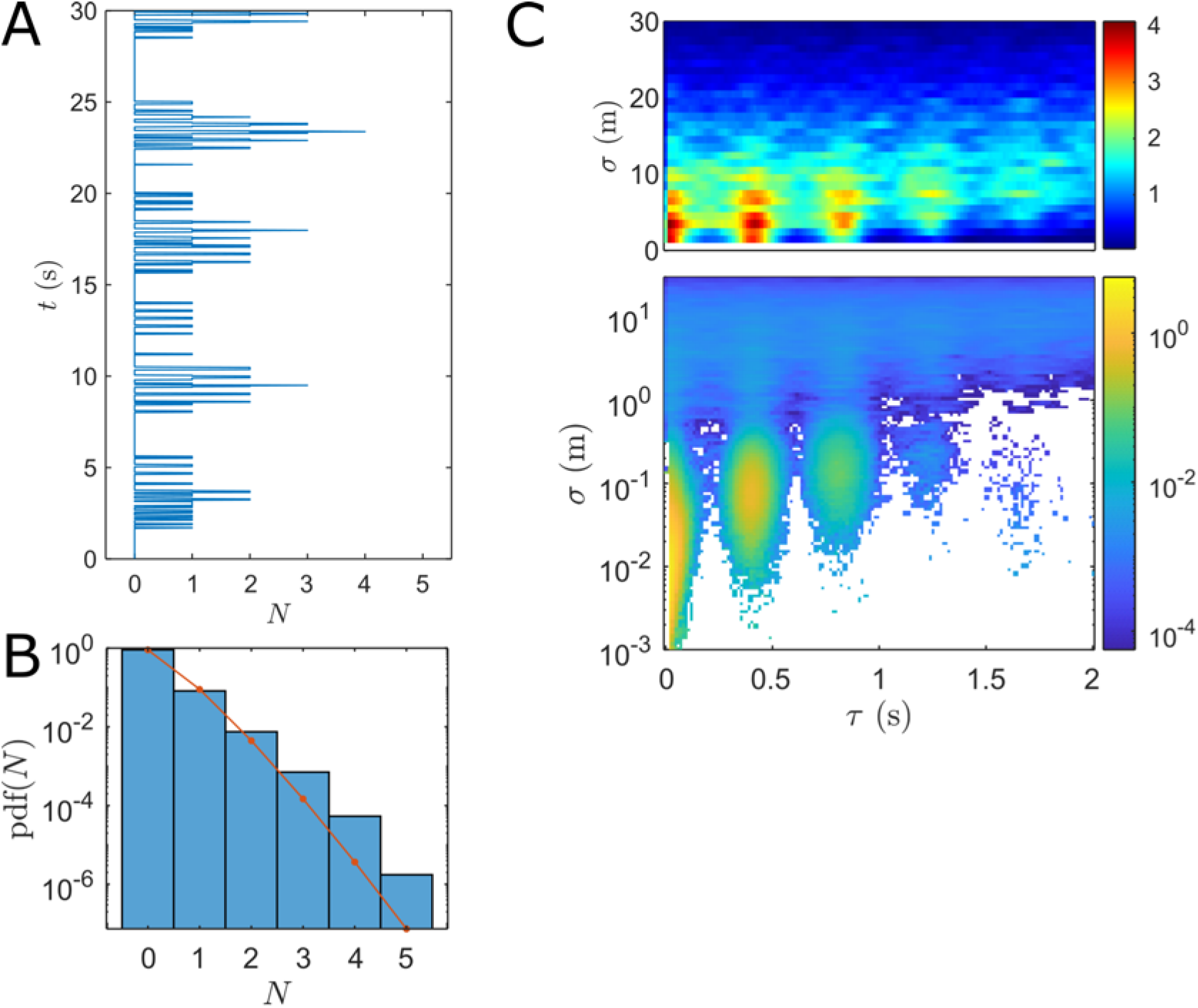
(A) Time series of the number *N* of flashes per frame (August 10). For visibility, only a short 30s-interval is shown. Many concurrent flashes are seen during repeated bursts of activity. (B) Experimental probability distribution (pdf) of *N* over 2hr of data. Red line is the result for a Poisson distribution with same average λ as pdf(*N*). (C) Spatiotemporal correlations: distribution of separation *σ* and time delay *τ* between flash occurrences. The bottom plot shows the full range of spatial separation, in logarithmic scale. The top plot shows the distribution when *σ* > 1m, emphasizing extrinsic correlations.

The validity of the Poisson distribution as the null hypothesis here should be nuanced. Indeed, flashes typically span not just one frame, but up to 5, and many are repeated three times within a short period. Therefore, the distribution of *N* contains internal correlations, even if flashers were fully independent. However, we show in the SI that persistent, repeated flashes in an independent correlation does not significantly alter the resulting distribution of test statistics from 1-flash-per-frame distributions, hence the comparison with the Poisson distribution still holds.

### 3.3 Spatiotemporal distributions reveal interactions

Synchronous behaviour was also made apparent from the distribution of spatiotemporal correlations between flash occurrences. For two 3D-reconstructed flashes of coordinates (*x_i_, y_i_, z_i_, t_i_*) and (*x_j_, y_j_, z_j_, t_j_*), we can calculate the corresponding separation *σ_ij_* = |(*x_j_, y_j_, z_j_*) – (*x_i_, y_i_, z_i_*)| and time delay *τ_ij_* = |*t_j_* – *t_i_*|. The 2D distribution of (*τ,σ*) is presented in Fig. 2C. Several features are significant. For *σ* < 1m, the correlations are narrowly distributed around three peaks at *τ* ≃ 0,0.45, 0.9s. These correspond to the flash triplets emitted by individual fireflies, which remain localized as they travelled typically no more than 0.5-1m over 1s. Due to the swarm’s low density, flashes from different fireflies happening within 1m from each other were very rare. Next, secondary peaks are visible for *σ* > 1m. These cannot originate from the same firefly, since they did not travel fast enough. They therefore represent extrinsic correlations. The fact that these peaks are narrow and distributed at the same specific times as intrinsic correlations is further proof of synchrony. Importantly, synchronizing respondents were always situated in the 1-10m range, as correlations vanish at larger distances. This confirms previous observations that synchronization information propagates only at short range [14].

## 4 Discussion and Conclusion

Remarkably, the collective flashing pattern of *P. knulli* appears to mirror that of another species of the same genus, *P. carolinus*. Both species happen to flash synchronously during bursts of activity lasting a few seconds and repeated periodically at high density, as evidenced by the frequency spectra of their respective time series (see SI, Fig. S4). Collective bursts are typically repeated every 12-14s for *P. carolinus*, but only every 5s for *P. knulli*. This difference seems most likely attributable to the individual flashing pattern, consisting of about three flashes for *P. knulli*, but six to eight for *P. carolinus*. Spatiotemporal correlations between flash occurrences further emphasize this distinction, but also a similar range in of distance interaction (see SI, Fig. S5). These qualitative similarities but quantitative differences offer an interesting opportunity to test mathematical models of intermittent synchrony.

Overall, our recordings largely confirm the observations made in Ref. [6], and provide data and statistical analysis to demonstrate the synchronous behaviour of *P. knulli*. While our video recordings confirm the existence of synchronization behaviour in this third North American species, the absence of enough firefly density prohibited the observations of more complex behaviours previously reported in Ref. [6]. It is clear that firefly activity in the Peña Blanca Canyon has declined significantly over the past few decades. The population decline is concerning [22]. While fireflies exist across the American West, populations tend to be much more sparse and localized than in the East. These populations are rare and fragile; *P. knulli* is now considered as Vulnerable by the International Union for Conservation of Nature [23]. Over the past few decades, increased habitat destruction (*e.g*., increased ranching and illegal ATV recreation) and changing weather patterns might be putting these species and populations at risk. While 2021 was an excellent monsoon season in Southern Arizona, prior years years had been excessively dry. These populations have also been less studied and less monitored than Eastern populations. Increased monitoring and research efforts seem important to further understand population dynamics. In the meantime, it seems crucial to encourage local initiatives to further protect firefly habitats.

## 5 Acknowledgments

For participation in the field and precious insights about the data, we would like to thank Larry Buschman, Lynn Faust, Cheryl Mollohan, Owen Martin, Andrew Moiseff, and Wesley Ryan.

## 6 Data Accessibility

Datasets of 3D reconstructed flash occurrences are made available at [TBD].

## 7 Author Contributions

R.S. performed experiments with guidance from J.M.C. and assistance from L.G. in the field. R.S. analysed the data. R.S. and O.P. wrote the paper, and all authors reviewed it before submission.

## 8 Conflict of Interest

The authors declare that they have no competing financial interests.

## 9 Funding

This work was supported by the BioFrontiers Institute at the University of Colorado Boulder.

## Supplementary Information

### 1 Goodness-of-fit tests for the Poisson distribution

The Poisson distribution of rate λ is defined as *Poiss*(*N* = *k*) = *e*^−λ^λ^*k*^/*k*!. We denote the first statistical moments by *μ*, *σ*^2^, *α, β* for mean, variance, skewness, and *excess* kurtosis (*i.e*., kurtosis minus 3), respectively. The empirical moments calculated from an experimental sample are written as 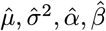. Theoretically, λ = *μ* = *σ*^2^ = *α*^−2^ = *β*^−1^. To test our empirical distributions of sample size *n* against the Poisson distribution, we used the two following test statistics:

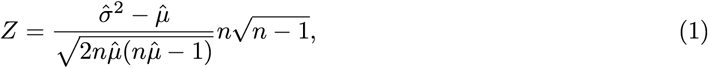

as defined in Ref. [1], and

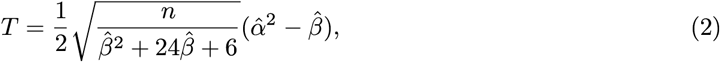

as defined in Ref. [2]. In the large *n* limit, these two test statistics converge in probability to the standard normal distribution (*μ* = 0, *σ*^2^ = 1). We indeed verify this convergence by numerical simulations of Poisson distributed samples of independent, one-frame flashes, see Fig. S1. Specifically, we simulated *N_s_* = 1000 “time series” of *n* = 10^5^ frames (same order of magnitude as our experimental time series), where each frame is independent from all others and contains a number of flashes drawn from a Poisson random generator with rates λ = 0.01,0.03,0.1, 0.3,1. The resulting distributions for *Z* and *T* are well approximated by the standard normal distribution; notably, their empirical moments are close to expected (Fig. S1C).

### 2 Simulation of the distribution of independent, but persistent and repeated, flashes

*P. knulli* flashes tend to persist for longer than the duration of a single frame (1/60s) and be repeated typically three times over an interval of 0.9-1s. Based on our videographic data, a typical flashing motif for an individual firefly is *M* = (1,1,0_20_,1,1,1,1,0_20_1,1,1,1). This is of course prone to variability in the duration of a flash (number of consecutive 1’s), interflash intervals (number *q* of zeros, 0_*q*_, between flashes), and overall number of flashes. But for simplicity we assume that all fireflies flash according to this pattern.

A better model for a swarm of *uncorrelated* fireflies, then, is to generate a Poisson distributed time series, but where each flash is convoluted with the motif *M*. The input rate λ is adjusted to match the desired output rate. Because flashes are persistent and repeated, this slightly increases the proportion of concurrent flashes. We calculate the *Z* and *T* statistics for these simulated time series (*N_s_* = 1000, *n* = 10^5^) with effective rates λ = 0.01,0.03,0.1, 0.3,1. The results are shown is Fig. S2.

The test statistics *Z* and *T* remain normally distributed around 0 (notably, their kurtoses are close to 0), but the repeated motif widens the distributions and increases the variance. For both *Z* and *T*, the new variances for λ ≃ 0.1 are approximately equal to 5 (Fig. S2C).

**Figure S1:**
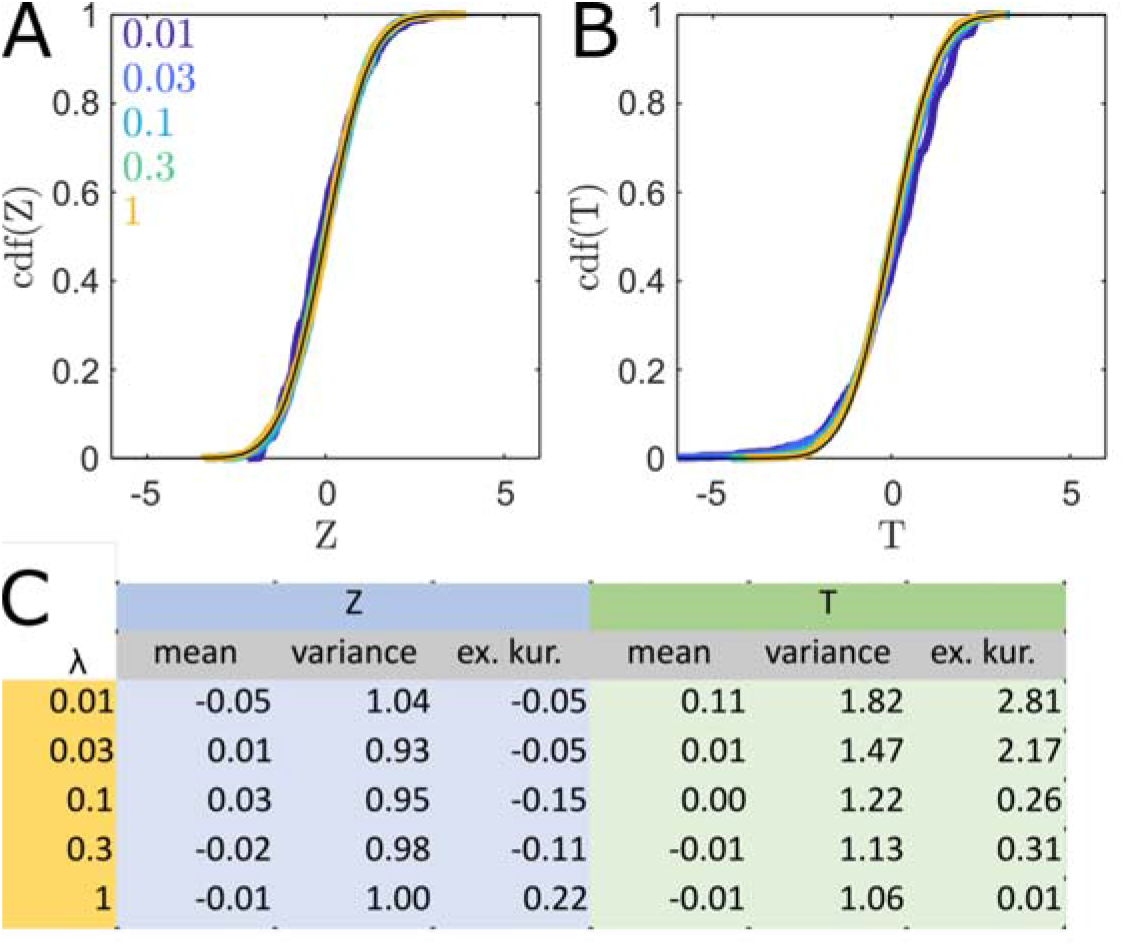
*Z* and *T* statistics for purely Poisson time series. (A) Empirical cumulative distributions for *Z* for different λ values which are indicated in the legend (0.01, 0.03, 0.1,0.3,1). The black curve is for the standard normal distribution. (B) Corresponding distributions for *T*. (C) Empirical moments for the *Z* and *T* distributions across λ. The standard normal distribution has mean 0, variance 1, and excess kurtosis 0.

### 3 Statistical evidence of synchrony across three datasets

Extensive stereoscopic video recordings were obtained for three different nights (August 7, 9, 10, 2021). Each recording lasted about 2hr 30min, and captured between 30,000 and 60,000 3D reconstructed flashes. The 10min-averaged flashing rate for each night is shown in Fig. S3A. These rates show some fluctuations, but remain overall comprised between 0.05-0.1 flash/frame. The distributions of *N* for each night are shown in Fig. S3B,C,D, along with the Poisson distributions with the same average flashing rate. In all cases, the proportion of large spikes (large *N* values) is significantly larger in the experimental data than expected for independent flashing events. All three datasets appear inconsistent with the Poisson distribution.

The August 10 dataset returned values *Z* = 55 and *T* = −28, as calculated from Eqs. 1,2. Since these test statistics are Gaussian, one can calculate the corresponding *p*-value as:

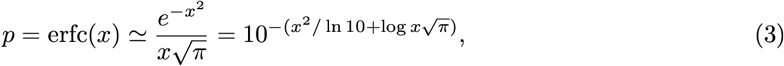

where erfc is the complementary error function. Using the increased variance for repeated flashes 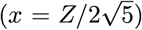, we find *p_Z_* ~ 10^−67^ and *p_T_* ~ 10^−18^. For August 9, *Z* = 54, *T* = −37, so *p_Z_* ~ 10^−65^ and *p_T_* ~ 10^−31^. For August 7, *Z* = 30, *T* = −20, so *p_Z_* ~ 10^−21^ and *p_T_* ~ 10^−10^. In all instances, the probability for synchronization is extremely high, within the proposed statistical model.

### 4 Comparison between collective flashing patterns of *P. knulli* and *P. carolinus*

The typical patterns of collective flashing in *P. knulli* (Pk) is compared to that of Eastern US species *P. carolinus* (Pc). Data about Pc was collected on June 10, 2020 in Great Smoky Mountains National Park, Tennessee (see Ref. [3] for details).

**Figure S2:**
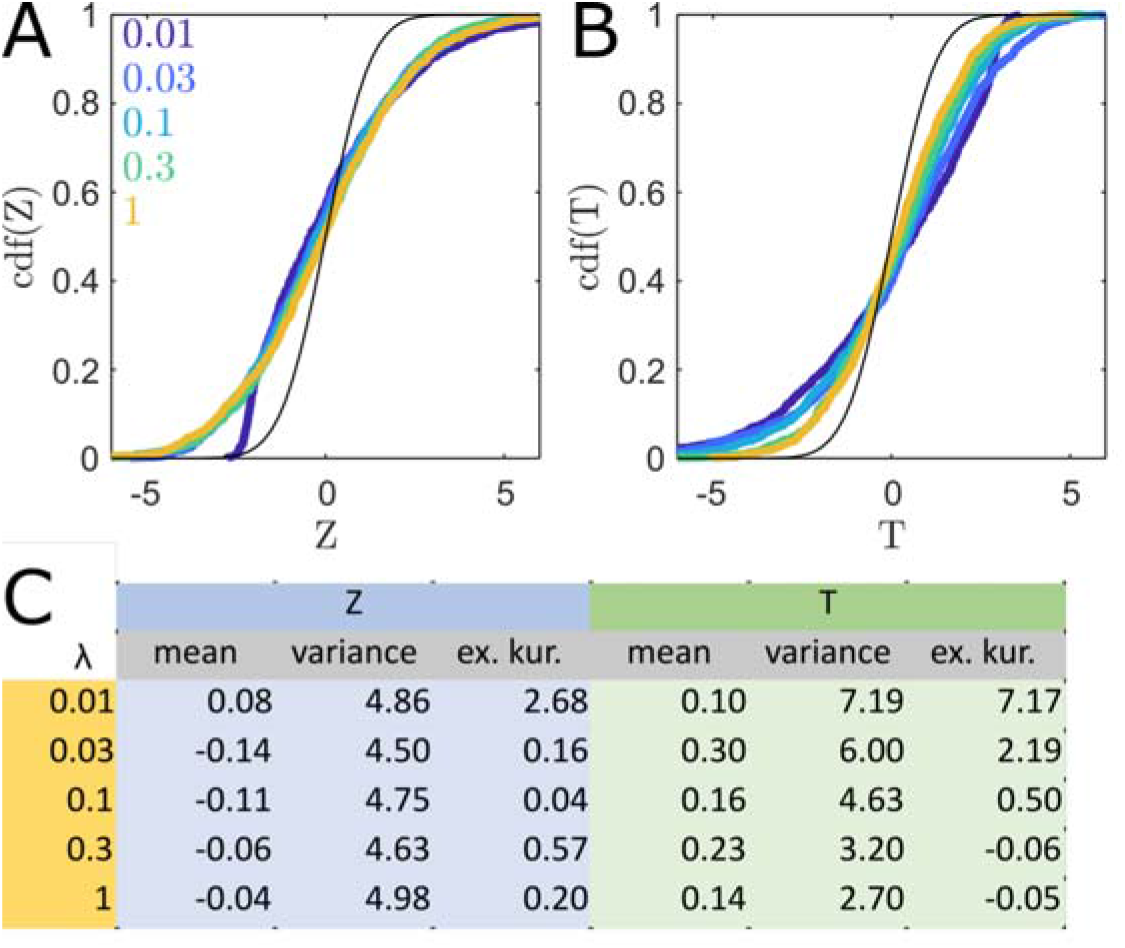
*Z* and *T* statistics for Poisson time series convoluted with flashing motif *M*. (A) Empirical cumulative distributions for *Z* values for different λ values which are indicated in the legend (0.01, 0.03,0.1,0.3,1). The black curve is for the standard normal distribution. (B) Corresponding distributions for *T* values. (C) Empirical moments for the *Z* and *T* distributions across λ.

#### 4.1 Temporal patterns

Both patterns consist of synchronous spikes of activity within periodic bursts (Fig. S5A). Calculating the Fourier transforms for each time series reveal two distinct underlying frequencies (see also Ref. [4]). The fast frequencies, about 2.0Hz for Pc and 2.4Hz for Pk, correspond to interflash intervals of 0.5s and 0.4s, respectively. In addition, a slow frequency, 0.09Hz (Pc) and 0.018Hz (Pk) correspond to burst periods of 11s and 5.5s, respectively. The structure of the temporal patterns are similar, but the values of the two main frequencies differ. Significantly, the average flashing numbers of Pc is significantly higher than for Pk, which also results in a higher signal-to-noise ratio (sharper spikes and more distinct bursts). This is in line with previous observations that showed that synchrony requires a large number of flashers, possibly because the flash information propagates locally.

#### 4.2 Spatiotemporal patterns

Similarly, the spatiotemporal correlations of both species share many similarities. At short length-scales (*σ* < 0.1 – 1m), self-correlations show flash trains for an individual firefly, typically three flashes for Pk and six for Pc. At larger length-scales (*σ* > 1m) extrinsic correlations are distributed along specific values of *τ* corresponding to the short periods of 0.4s and 0.5s uncovered in the frequency spectra as well. These indicate flash synchronization. Again, the peaks are sharper in the case of Pc, due to the swarm’s higher density.

**Figure S3:**
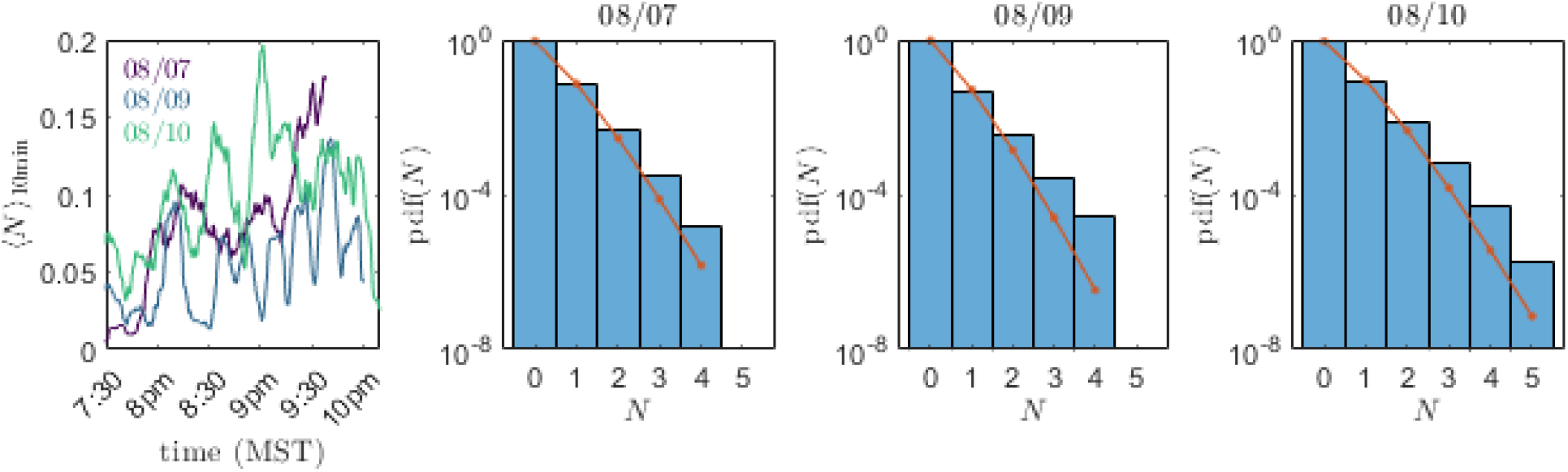
(A) Moving averages (10min, or 36000 frames) of *N* time series for the three recordings. (B-D) Experimental distribution of *N* for all three nights (blue histogram), and Poisson distribution with the same average flashing rate (red line).

**Figure S4:**
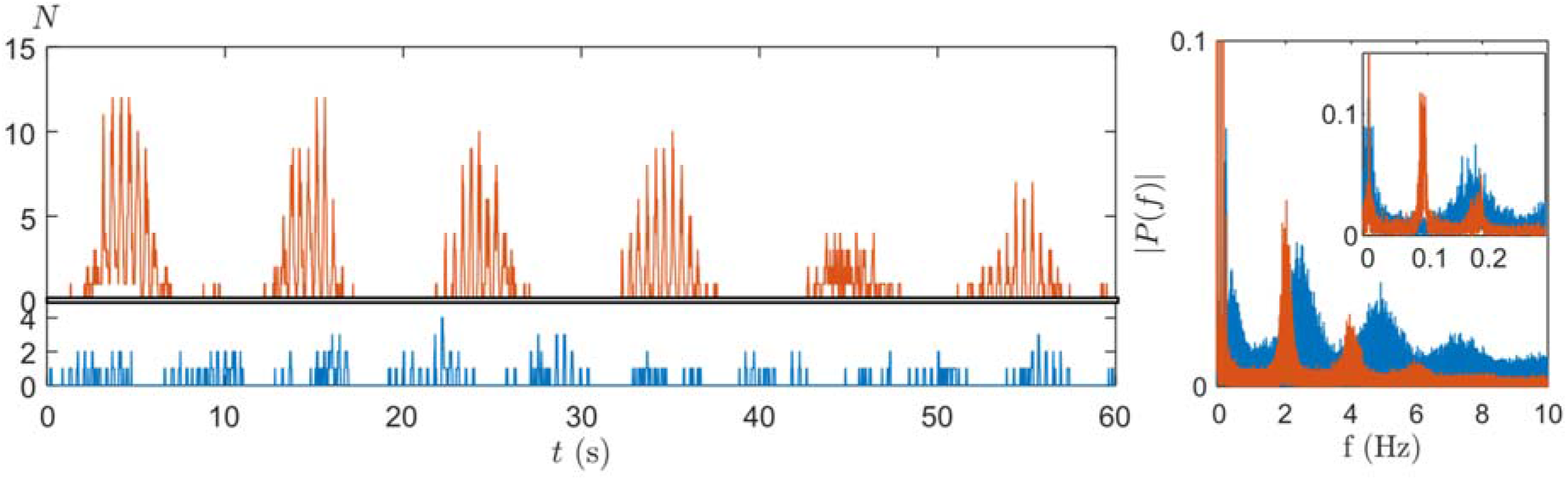
(A) Time series of *N* for *P. knulli* (blue) and *P. carolinus* (red). (B) Corresponding Fourier Transforms, revealing the main frequencies. The inset shows the frequency spectra a low frequencies (large periods).

**Figure S5:**
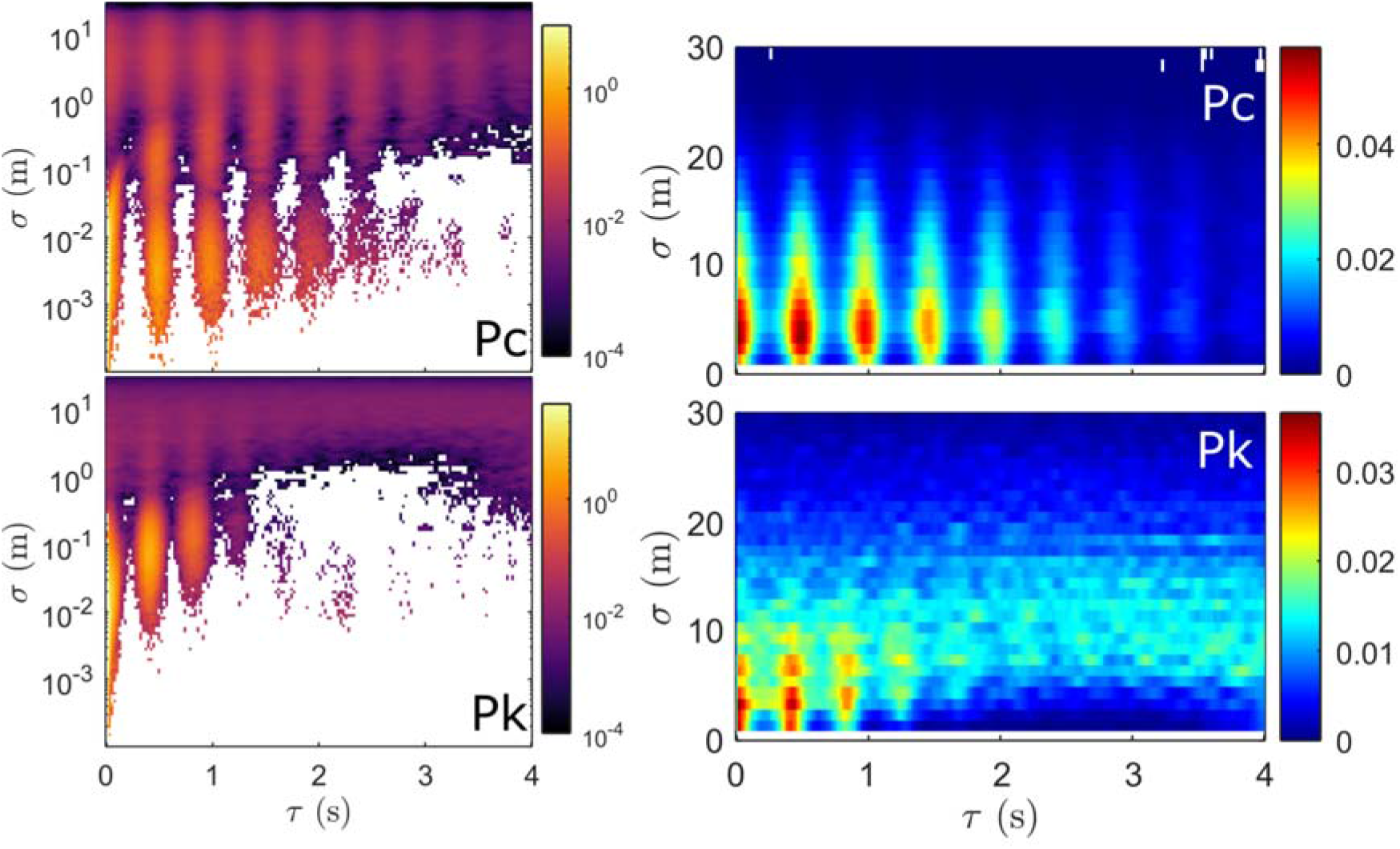
Comparison of spatiotemporal correlations between both species (Pc and Pk). (Left) Distributions across the full range of distances *σ* = 1mm - 30m in logarithmic scale. (Right) Distributions for *σ* > 1m.

## Notes

### Competing Interest Statement

The authors have declared no competing interest.

